# Quality assessment of tissue samples stored in a specialized human lung biobank

**DOI:** 10.1101/407411

**Authors:** M. Lindner, A. Morresi-Hauf, A. Stowasser, A. Hapfelmeier, R. Hatz, I. Koch

**Affiliations:** Asklepios Biobank for Lung Diseases, Department of Thoracic Surgery, Asklepios Fachkliniken München-Gauting, 82131 Gauting, Germany; Member of the German Center for Lung Research – DZL; Asklepios Biobank for Lung Diseases, Department of Pathogy, Asklepios Fachkliniken München-Gauting, 82131 Gauting, Germany; Member of the German Center for Lung Research – DZL; Institute of Medical Informatics, Statistics and Epidemiology, Klinikum rechts der Isar, Technical University Munich, 81675 Munich, Germany

## Abstract

Human sample, from patients or healthy donors, are a valuable link between basic research and clinic. Especially in translational research, they play an essential role in understanding development and progression of diseases as well as in developing new diagnostic and therapeutic tools. Stored in biobanks, fast access to appropriate material becomes possible. However, biobanking in a clinical context faces several challenges. In practice, collecting samples during clinical routine does not allow to strictly adhere to protocols of sample collection in all aspects. This may influence sample quality to variable degrees. Time from sample draw to asservation is a variable factor, and influences of prolonged storage at ambient temperature of tissues are not well understood. We investigated whether delays between 5 minutes and 3 hours, and the use of RNAlater RNA-preserving reagent would lead to a relevant drop in sample quality, measured by quantitative mRNA expression analysis. Our findings suggest that even under ambient conditions, delays up to 3 hours do not have a major impact on sample quality as long as the tissue remains intact.

## Introduction

In 2008, a biobank was founded at the Asklepios Clinics in Gauting, a clinic specialized on thoracic diseases. By the end of 2017, it contained solid tissue and liquid biomaterials from nearly 4000 patients, up to 45.000 aliquots. 12.000 of these are aliquots of solid tissue samples of tumor and peritumor tissue from patients suffering from various bronchial carcinomas, lung metastasis of other types of cancer and tissue from benign thoracic malignancies. Serum, plasma, BALF fluids, cell pellets and pleural effusions are collected, whenever possible as paired samples. The Biobank is integrated into the German Center for Lung Research (DZL). Biobanks at all sites of the DZL aim to collect samples according to harmonized Standard Operation Procedures, making samples comparable among the sites and their usage in common scientific projects reliable.

After obtaining patients‘ broad informed consent, based on the suggestions of the German Ethics Council, samples are collected during routine clinical procedures. With the exception of blood specimens, only clinical remains are stored. Tissues are collected after diagnostic procedures have been completed. Some effects affecting sample quality cannot be influenced, like warm ischemic time (i.e. the time between truncation of the blood supply and removal of the tissue from the body), others can be controlled more or less satisfactorily. Delay between withdrawal from the body and asservation is kept as short as possible, but is subject to variation due to clinical routines. This means, that standardization of sample collection is a challenge, which cannot regularly be met in everyday clinical life to a full extent. Nevertheless, the aim is to gain samples of highest quality, suitable for a variety of scientific questions and methods including molecular biological analyses like expression profiling.

Collection of solid samples is often done by snap freezing in liquid nitrogen, followed by long-term storage at -80 °C or in liquid nitrogen. This guarantees conservation of biological processes at the moment of freezing, however is difficult to integrate into the clinical routine. Alternatives such as incubating samples in protecting reagents like RNAlater^®^ or ProtectAll^®^ before freezing are gaining more and more importance(1-3). This study aims to compare snap freezing and incubation in an RNA stabilizing reagent with regard to sample stability, nucleic acid recovery and reproducibility of mRNA expression measurements. Influences of pre-freezing delays are also addressed.

## Materials and Methods

The Asklepios Biobank for Lung Diseases was approved by the Ethical Committee of the Ludwig-Maximilians-University of Munich in February 2011 (Project-No. 330-10)

### Tissue collection

Tissue samples after operative procedures are routinely collected by a pathologist in parallel to diagnostic procedures. Samples not required for patients‘ diagnosis are made accessible to the biobank. In a standardized fashion, samples from 4 patients were processed after 5 (±2), 20 (±5), 60 (±10), and 180 (±10) minutes of cold ischemic time to mimic variability in clinical sampling. Patients’ characteristics are summarized in table 1. At each time point, pieces cut to a maximum size of 5×5×5 mm were either snap frozen in liquid nitrogen before storing at -80 °C, or transferred to RNAlater™ (Quiagen, Hilden, Germany). RNAlater samples were left at 4 °C for 24 h or 7 days, before the RNAlater was discarded and samples were stored dry at -80 °C without snap freezing. Thus, a set of 12 samples (4 time points and 3 methods) was generated for each patient. (Fig 1).

**Table 1:** Patients’ characteristics

| Patient No | Sex | Age | Diagnosis | Stage |
| --- | --- | --- | --- | --- |
| 1 | male | 74 | Large cell carcinoma neuroendocrine | Ib |
| 2 | male | 73 | Adenocarcinoma | IIb |
| 3 | female | 53 | Squamous cell carcinoma | IIa |
| 4 | female | 54 | Typical carcinoid | Ib |

**Fig. 1.**
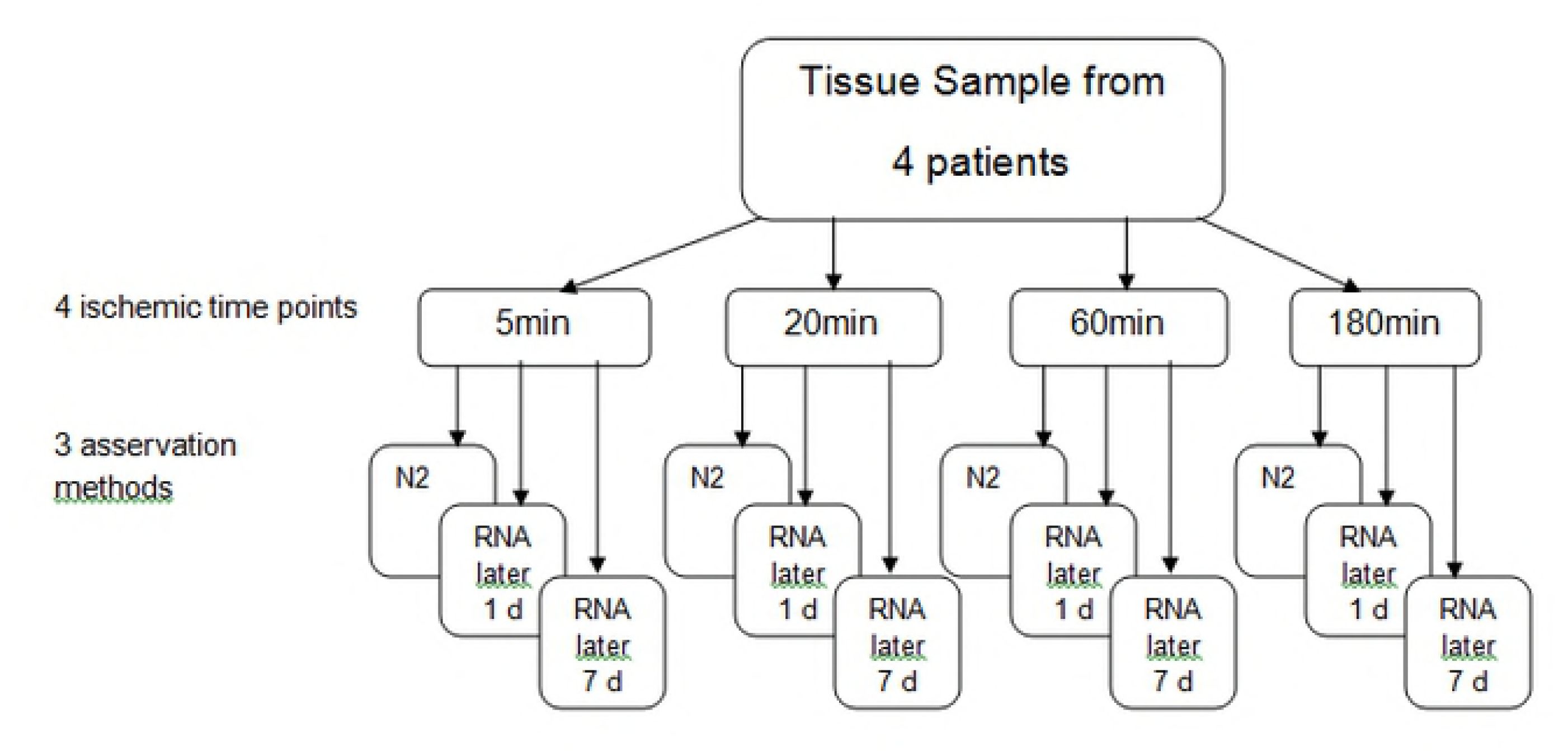
Experimental design. Tumor tissue samples from 4 lung cancer patients were cut into aliquots and processed after 4 different periods of time, with 3 different methods before long term storage and subsequent RNA isolation.

### Sample Processing, nucleic acid extraction and cDNA synthesis

Frozen samples were mounted on a precooled object plate for cutting in a NX70 microtome (Thermo Fisher Scientific, Waltham, Massachusetts) at -30°C, using MX35 tempered microtome blades for hard tissue (Fisher Scientific, Waltham, Massachusetts). After trimming, a 4 µm section was cut for HE staining and pathological evaluation. 5 x 10 µm sections were cut for RNA isolation, followed by another 4 µm HE section, before 5 x 10 µm section were cut for DNA isolation and a final 4 µm HE section.

RNA and DNA were isolated using RNeasy Mini Kit or QiaAmp Mini Kit (Qiagen, Hilden, Germany) respectively, according to the manufacturer’s instructions. Quantity and quality of nucleic acids was assessed by measuring OD at 260/280 nm in a P330 nano photometer (Implen, Munich, Germany) and determination of RIN values (only RNA) using the Agilent 2100 BioAnalyser™ (Agilent Technologies, Waldborn, Germany). 1 µg of total RNA was reversed transcribed into cDNA using random hexamer primers and superscript II reverse transcriptase (Life Technologies / ThemoFisher Scientific, Waltham, Massachusetts).

### Real-Time qPCR

RT-qPCR Assays for EGFR, ERCC1, RRM1 and HIF1 and TBP as a so called housekeeping gene were performed on LC480 light cycler (Roche, Mannheim, Germany), either using a SYBR-Green assay (MesaBlue, Eurogentec, Liège, Belgium), or light cycler FRET probes from Roche’s universal probe library. Primers and probes and cycling parameters are depicted in table 2. 18S rRNA was used to normalize expression levels. Expression values were expressed as absolute expression levels (copies/18S) for each gene, or as relative expression levels to the average expression measured in the 12-sample set of each patient to make intersample differences easily comparable among different genes, regardless of their absolute expression levels.

**Table 2:**
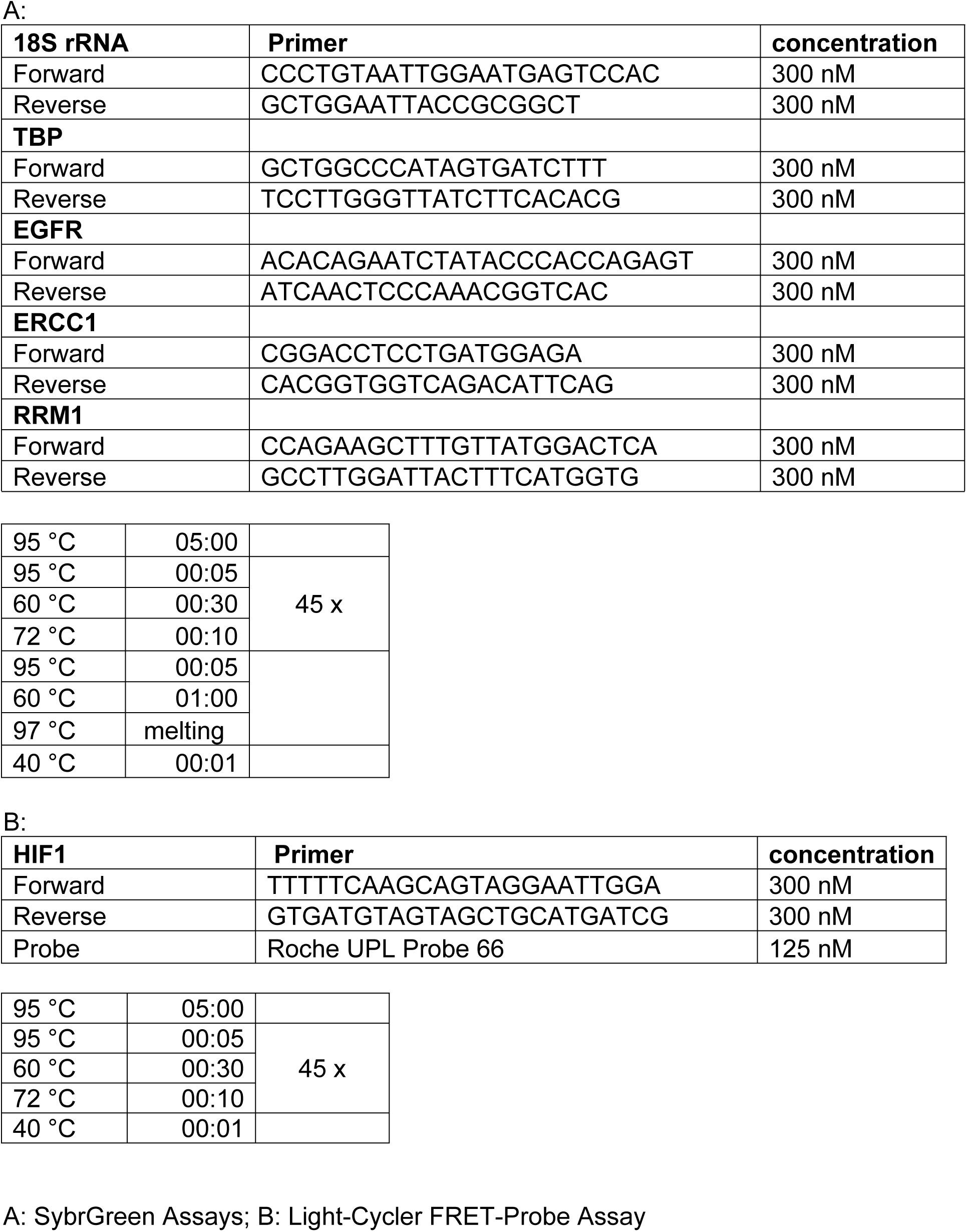
Real-time qRT-PCR Assays

### Statistical analysis

Statistical analysis was performed using IBM SPSS Statistics for Windows, version 23 (IBM Corp., Armonk, N.Y., USA) and R 3.4.2 (R Foundation for Statistical Computing, Vienna, Austria). The distribution of qualitative and quantitative variables is described by absolute and relative frequencies and means ± standard deviation, respectively. Repeated measures ANOVA was used for hypothesis testing of differences in asservation methods and time points. Bland-Altman analyses were performed by the alternating regressions approach to account for repeated measurements and assuming a constant bias in the conversion of methods (4). All statistical tests were conducted on two-sided, exploratory 5% significance levels.

## Results

Except for the standardized delay times, sample collection fully resembled the routine collection procedure. Delay times were chosen to mimic the intersample variability regularly imposed onto the samples on their way from the operation room to pathology and finally to the biobank, whereat a delay time of less than 5 minutes is essentially never met during daily collection. Most of the samples reach the biobank within 20 minutes of cold ischemic time, and all samples with cold ischemic times of more than 60 minutes are usually discarded.

### Histological evaluation

Pathological examination is critical before using archived human material. We compared HE-stained slices of fresh frozen or RNAlater preserved tissue. To be able to cut tissue samples treated with RNAlater, it turned out to be necessary to cool the cryostat to -30 °C to ensure the sample kept frozen, and to use tempered blades normally used to cut hard tissues like bone. Doing so, we were able to cut these tissues without removing RNAlater. Both – tissue that was snap frozen and tissue preserved in RNAlater, were equally well suited for histological evaluation (Fig. 2)

**Fig. 2.**
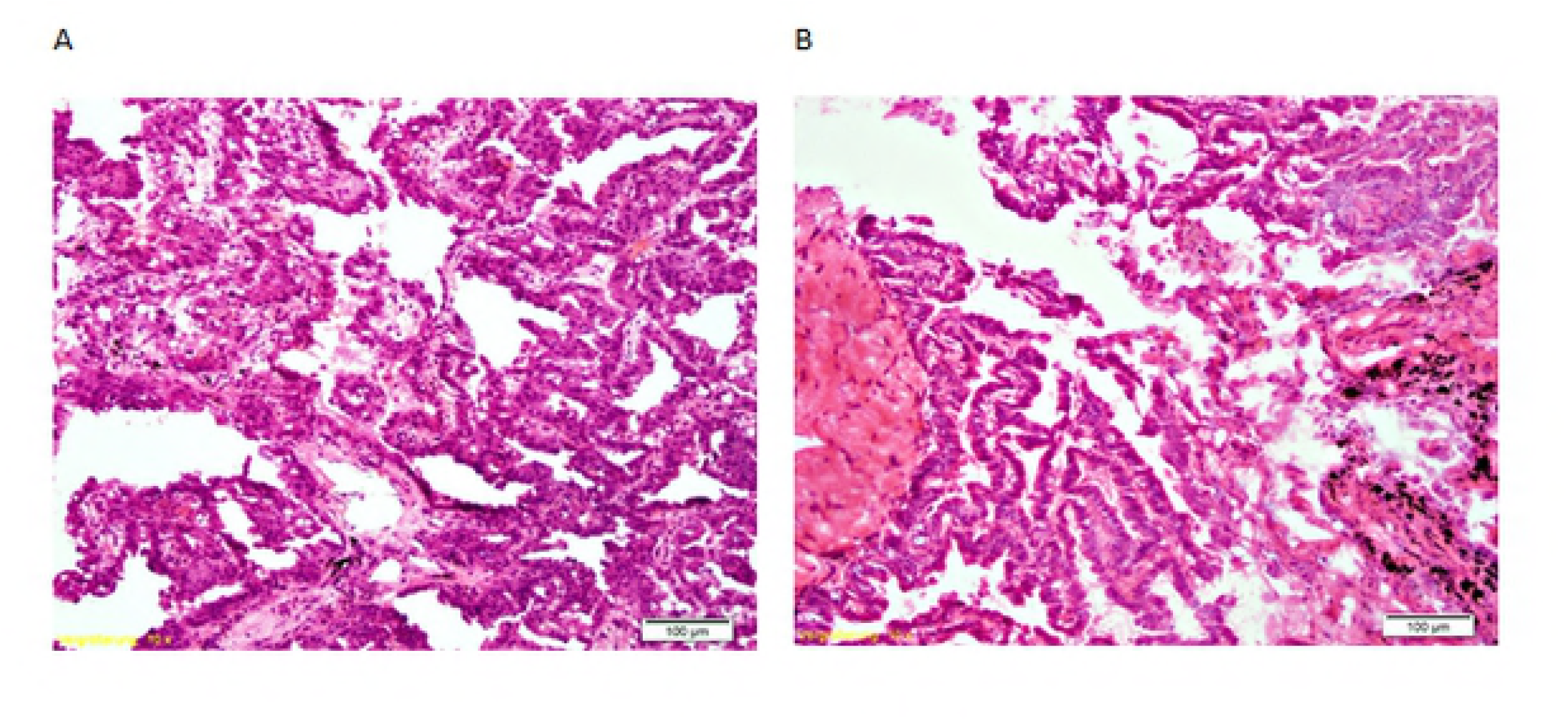
Histological evaluation. HE stained histological section of adenocarcinoma, patient 2. A: tissue preserved in RNAlater for 1 day; B: tissue snap frozen in liquid N_2_

### Influence of asservation method and cold ischemic times on nucleic acid quantity and quality

Due to the heterogeneity of the tissue, the quantity of nucleic acids isolated from the tissue specimens is influenced by a variety of factors such as tissue size, cell number, percentage of necrosis etc.. It is assumed that DNA would be stable under all conditions tested. RNA quantity relative to DNA quantity was also rather stable, regardless of shorter or longer ischemic times (data not shown). With regard to RNA quality, we measured the RIN values (RNA integrity number) for all samples. RIN values showed negligible variation. 60 % (29) of all samples had RIN values of 9 or higher, 27 % (13) of 8-9, 10 % (5) of 7-8 and 2 % (1) of lower than 7, with a mean of 9.07 (SD 0.86). There was only a minor difference between mean RIN values of fresh frozen tissues samples and RNAlater preserved ones (9.04, SD 0.86) vs. 9.08, SD 0.86, Fig.3.).

**Fig. 3.**
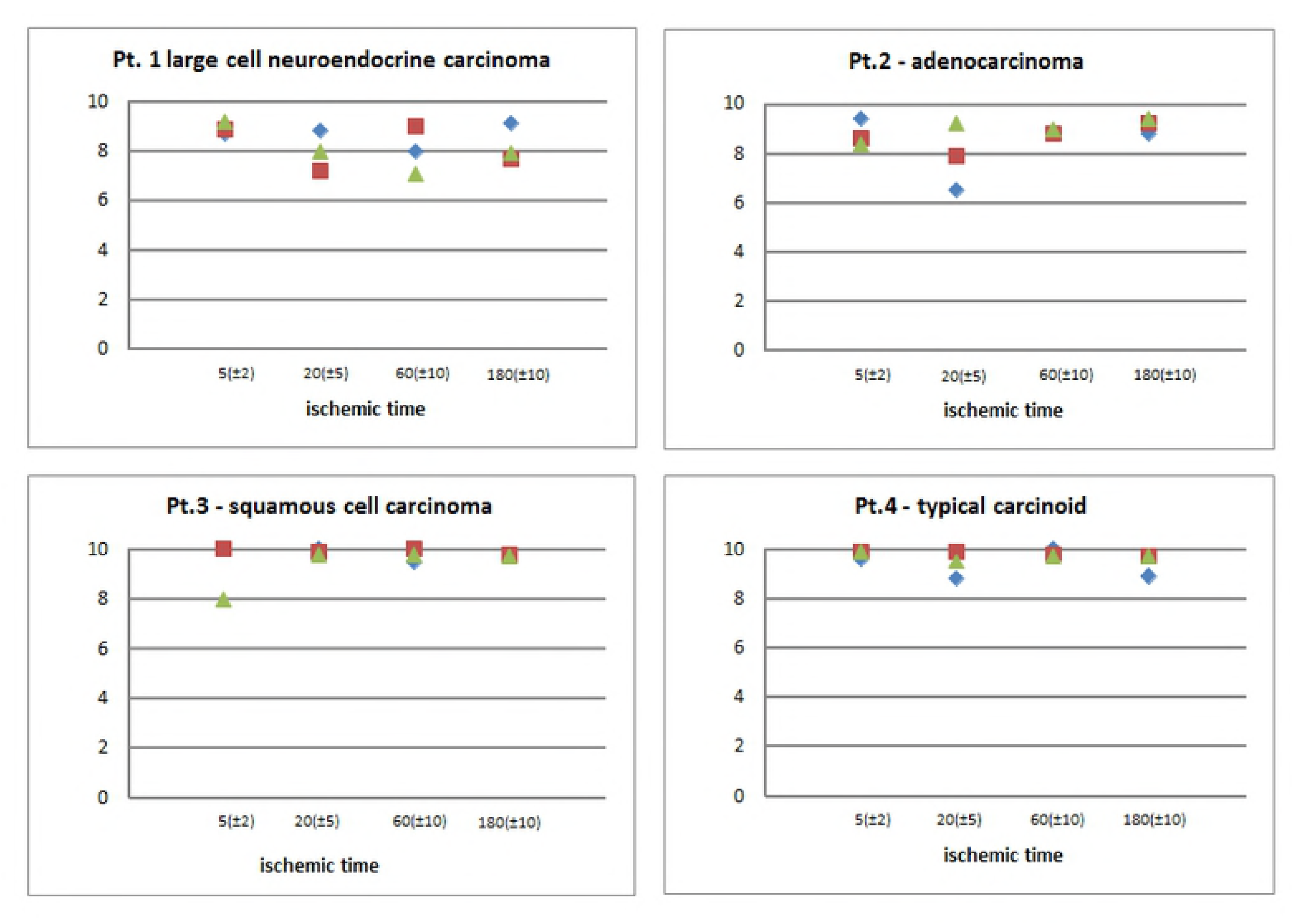
Influence of cold ischemic time and asservation procedure on RNA quality. Overall RNA quality isolated was assessed on an Agilent BioAnalyser. RIN (RNA integrity numbers) were compared between different ischemic times and different asservation methods (blue: shock frozen in liquid N_2_; red: RNAlater, 1 day 4 °C; green: RNAlater, 7 days 4°C; all samples were stored at -80°C thereafter.

RIN-values – whether from tissue preserved in RNAlater or fresh frozen in liquid nitrogen - were not notably influenced by prolonged ischemic times up to 3 hours. Mean RIN values were 9.18 (SD 0.66) for an ischemic delay of 5 minutes, 8.79 (SD 1.11) for 20 minutes, 9.13 (SD 0.84) for 60 minutes and 9.14 (SD 0.69) for 180 minutes.

### Influence of asservation method and cold ischemic times on gene expression

RIN values only give a global impression of RNA integrity, precisely of the integrity of 18S and 28S rRNA. mRNAs might be prone to more or less rapid degradation or changes in expression profile. We therefore measured the expression levels of 5 mRNAs by qRT-PCR on a LC480 light cycler device. All results were normalized to the content of 18S rRNA, as measured by qRT-PCR. The genes tested were TBP as a housekeeping gene, EGFR, ERCC1 and RRM1 as genes with potential predictive roles for lung cancer therapy, and HIF1 as a gene regulated by hypoxia, at least partially on RNA level. Reliable results were obtained with all RNA samples. As expected, expression levels varied between the four patients to a great extent. To compare gene expression between patients, we used the mean measured values of all 12 tested samples (Fig. 4). Mean expression values of the housekeeping gene TBP varied by a factor of 2 between individual patients. The maximal interindividual difference was 40 % for ERCC1, 60 % for RRM1 and 280 % for HIF1. A 23fold variation was found for EGFR, with the squamous cell carcinoma sample showing the highest expression.

**Fig. 4.**
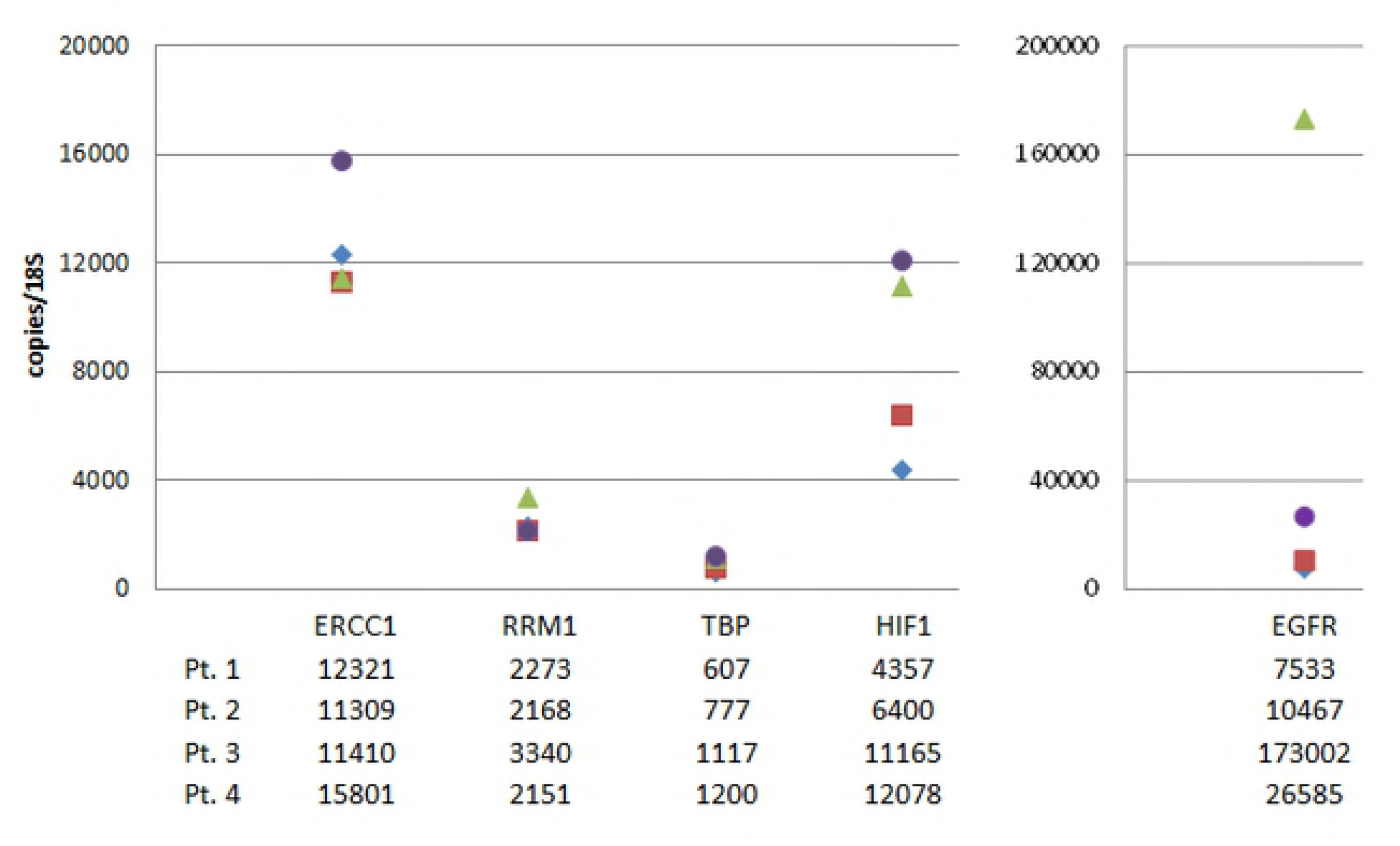
Expression levels of 5 mRNAs in 4 patients. Copy numbers were determined by qRT-PCR, normalized to 18S rRNA content as measured by qRT-PCR. Mean values of 12 aliquots are presented for each patient (blue: patient 1; red: patient 2; green: patient 3; purple: patient4).

Intraindividually, between the samples of one patient, variability was mostly within the range of a factor of 2, and ranging up to a factor of 5 in rare cases. There was no general trend towards lower mRNA content for longer ischemic delay, or for one of the asservation methods. None of the genes seems to be more susceptible to degradation within 3 hours after sampling. Fig. 5 exemplarily shows the results for HIF1, demonstrating the arbitrary distribution of variation.

**Fig. 5.**
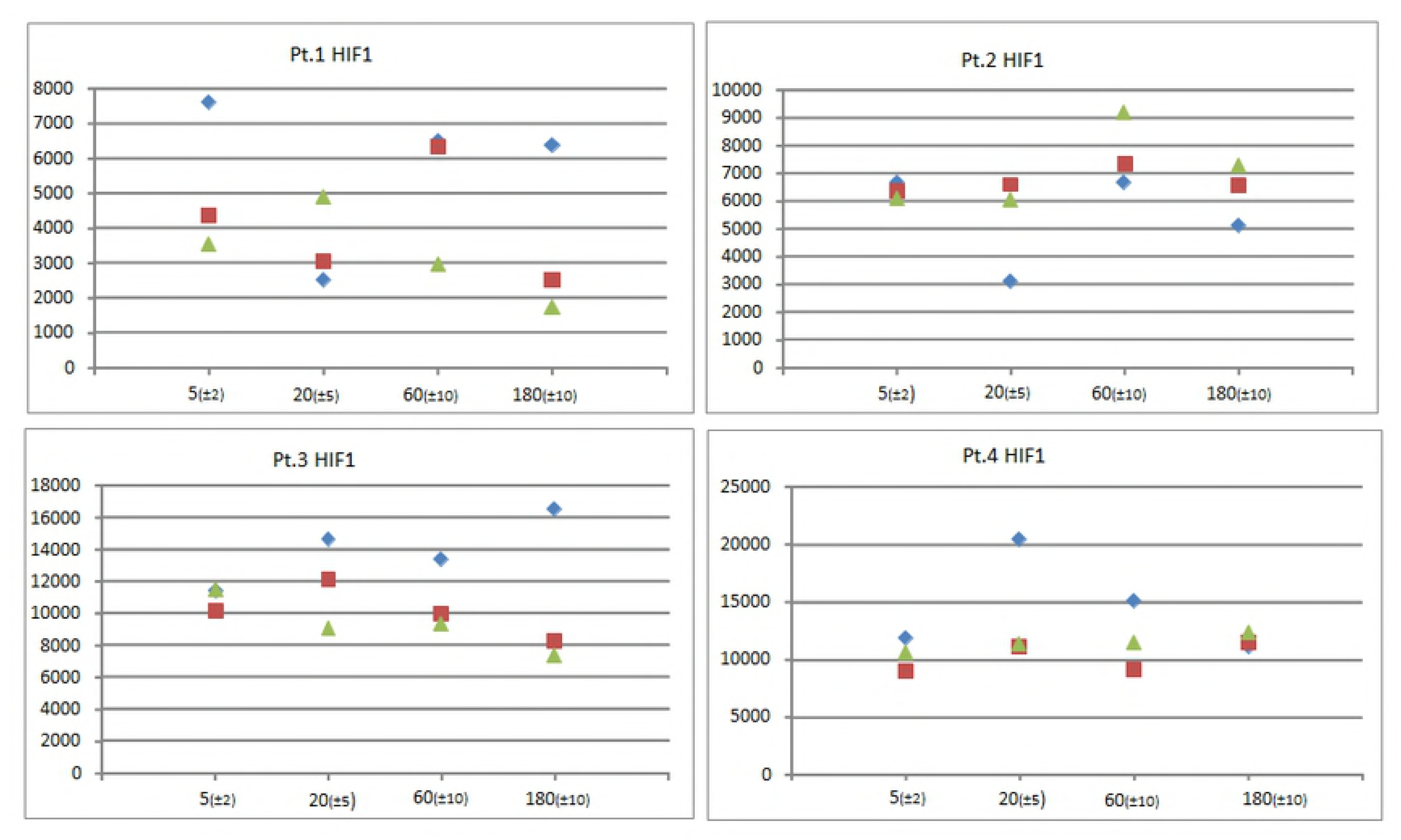
Expression of HIF1 mRNA. Expression was measured by qRT-PCR in four individual patients. Samples were either fresh frozen in liquid nitrogen (blue), or preserved in RNAlater for 1 day (red) or 7 days (green) before long term storage at -80 °C.

As expected, the greatest source of variation is introduced by interpatient differences in expression, superposing the influence of all other variables. In order to draw a more general conclusion about the comparability of the asservation methods, we normalized the data for each individual patient and each gene separately to the mean values of all 12 samples for this patient. Doing so, absolute expression levels no longer influence further analysis, and data oscillate around 1, with relative variation unaffected. It thus became possible to match genes with great differences in expression levels. A repeated measures ANOVA was used for hypothesis testing on differences between the asservation methods (p = 0.450) and the different ischemic times (p = 0.963). The variation for the average normalized expression values within the set of the 12 samples per patient for individual genes did not show preferences for a method or shorter ischemic times (Table 3). Scatter plots as well show no evidence for a trend in ischemic time (Fig. 6).

**Table 3:** Variation for average gene expression. Variation for the average normalized gene expression values for individual genes with regard to ischemic time and asservation method as assessed by repeated measure ANOVA.

| Time points | Mean | Std. Error | 95% Confidence Interval |  |
| --- | --- | --- | --- | --- |
|  |  |  | Lower Bound | Upper Bound |
| 5(±2) | 1.010 | 0.083 | 0.745 | 1.275 |
| 20(±5) | 0.997 | 0.085 | 0.728 | 1.266 |
| 60(±10) | 1.030 | 0.056 | 0.853 | 1.207 |
| 180(±10) | 0.963 | 0.095 | 0.659 | 1.267 |

| Methods | Mean | Std. Error | 95% Confidence Interval |  |
| --- | --- | --- | --- | --- |
|  |  |  | Lower Bound | Upper Bound |
| N2 | 1.100 | 0.093 | 0.805 | 1.395 |
| RNAlater 1d | 0.962 | 0.074 | 0.726 | 1.199 |
| RNAlater 7d | 0.937 | 0.052 | 0.773 | 1.102 |

**Fig. 6.**
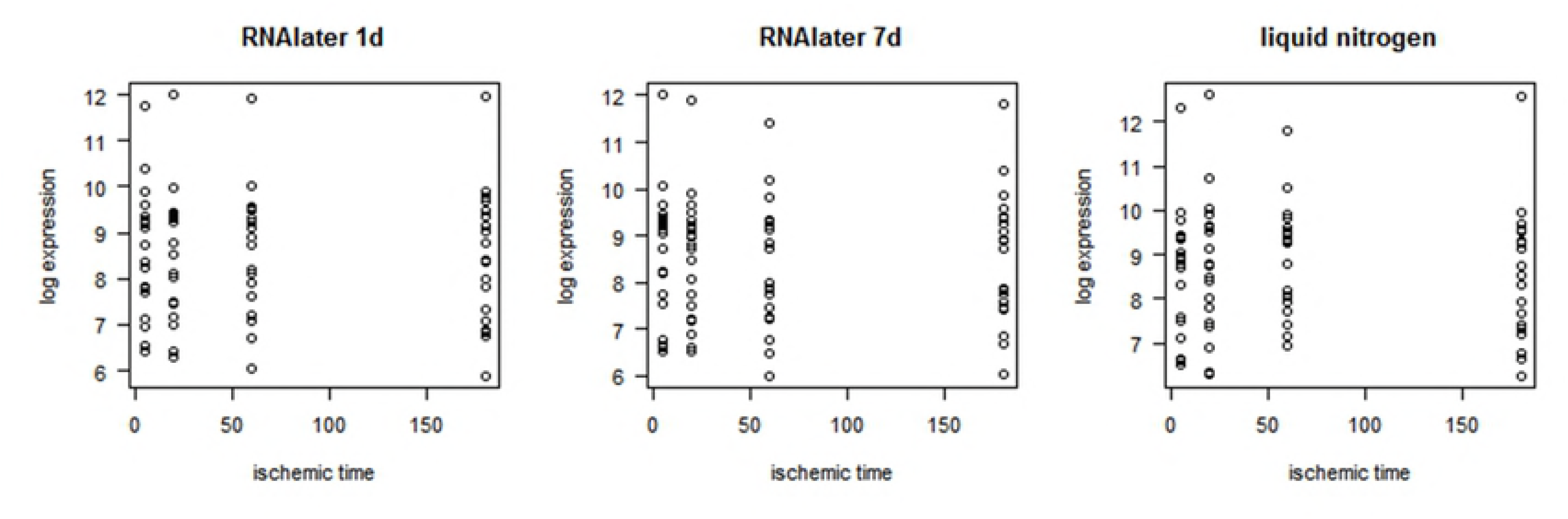
Variability between asservation methods and ischemic delay times. Mean normalized expression values of five selected genes. Individual expression levels were normalized to means for each gene and patient and then averaged over all patients for four different ischemic time points. Asservation methods: blue: liquid N_2_; red: RNAlater, 1day; green: RNAlater, 7d

To further evaluate the conformance of the methods, we performed Bland-Altman analyses (Fig. 7). Agreement between RNAlater RNA-preserving methods is slightly stronger (95% of expected relative deviations lie between 0.48 and 2.00) than between any of the RNA-later preserving methods and the liquid nitrogen snap freezing method (95% of expected relative deviations lie between 0.33 and 2.38), but there is no evident trend favoring one or the other method.

**Fig. 7.**
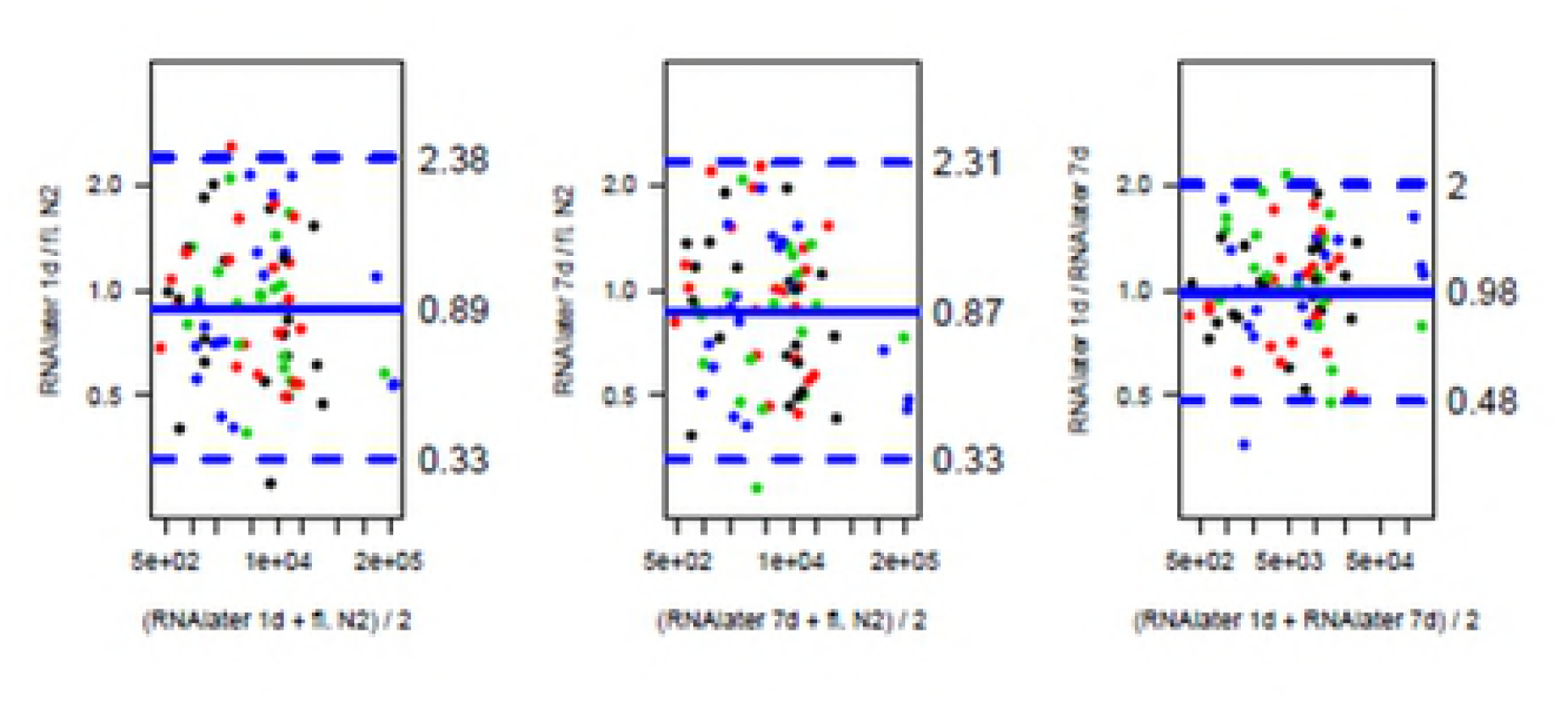
Scatterblots of mRNA expression values. Samples were processed in RNAlater (1 or 7 days) or by snap freezing in liquid nitrogen. Assuming a log-normal distriburtion of expression values, all computations were performed on a log scale.

**Fig 8: Conformance of RNA processing** Bland-Altman Plot demonstrating the conformance of RNA processing in RNAlater for 24 h or 7 days or snap freezing in liquid N2. The colours represent the different ischemic times (black 5’, red 20’, green 60’, blue 180’).

## Discussion

Biobanks constitute research infrastructures, providing samples for a wide variety of scientific purposes and methods. Randomly collected samples are compiled into cohorts with specific features such as diseases or therapies. It is of outstanding importance, that during the procedure of sample collection, processing and storage, the inherent characteristics of the samples are preserved, while introducing as little alterations as possible. Immediate stabilization of the expression pattern is a prerequisite if samples are to be used in mRNA expression analysis later.

Several authors could demonstrate good overall DNA and RNA stability and preservation of the global expression profile has been demonstrated in various ex vivo or post mortem conditions (1-3). Nevertheless, in tissues, briefly after harvesting, changes in the mRNA expression pattern are suspected to and have been shown to occur in a fraction of mRNAs, and is more pronounced if the tissue is exposed to room temperature rather than kept on ice (1, 3). This may be due to degradation processes, but also due to transcriptional changes induced by the altered environment, like lack of oxygen supply, and may vary in different types of tissue.

The gold standard to ensure preservation of the in vivo expression profile is to collect samples in a very standardized way, and to keep the time from harvesting to freezing as short as possible. Clinical studies or population based efforts of biobanking, collecting samples in the study context, adhere to strict standard operation procedures for sample acquisition. Sample collection in clinical context follows a different concept of biobanking. At the time of collection there is no specific scientific question or no specific sample characteristic to define inclusion criteria, however, samples are collected during routine clinical procedures, at various sites of the clinic, and often as remains after completion of several diagnostic procedures. A 100 % standardization of preanalytic procedures is thus impossible.

There are several ways to minimized intersample variability: keeping time from sample acquisition to freezing as short as possible, standardizing samples handling once arrived in the biobank’s lab and the use of protective reagents such as RNAlater, among others.

RNAlater is a high salt ammonium sulfate aqueous solution specified to stabilize RNA in solid tissue by precipitating out RNAses in a concentration and ph-dependent manner, which has been describe as early as 1974 (5).

Several studies have demonstrated good preservation of expression profiles in RNAlater preserved vs shock frozen tissue. This holds true for selected genes as measured by real-time PCR (6-9), and for RNA expression microarray analysis (10-12). DNA suitable for PCR analysis can be extracted from RNAlater stabilized tissue, but it remains uncertain whether protein analysis is possible in these samples. It has been shown, that quantitative proteomic analysis yields comparable results for snap frozen and RNAlater preserved biopsies of colon mucosa (13). Preliminary experiments using protein extracts of NSCL cancer biopsies in Western Blots suggest, that the protein and phospho-protein analysis will also be possible for selected proteins, but it remains to be demonstrated that this is generally conferrable (data not shown). Further experiments will be needed before the widespread use of RNAlater samples for protein or proteomic research.

Another preserving reagent, ProtectAll, will stabilize RNA, DNA and proteins. However, it has a major disadvantage for clinical biobanking. While RNAlater preserved tissue samples can be cut on a microtome and stained for histological analysis, this is not possible with ProtectAll preserved samples, since even at -80 °C ProtectAll will not be frozen but rather remain gelatinous and therefore samples cannot be cut. Sample characterization with respect to verification of diagnosis, tumor cell content etc. is thus not directly possible in these samples.

Using RNAlater during routine solid sample collection in various sites in the clinic has some major advantages: dangerous and expensive handling of liquid nitrogen can be omitted, samples can be placed directly into the preserving agent at the site of extraction by any clinician and be processed later in the biobank’s lab, and nucleic acids, in particular RNAs, are stabilized. On the other hand, it has to be taken into account that preservation is not as abrupt as shock freezing in liquid nitrogen, since RNAlater or any other similar reagent will need to diffuse into the samples. Samples have to be incubated in RNAlater for at least 24 h to ensure sufficient absorption before freezing, following the manufacturer’s instructions. During this time, RNAlater will salt out proteins such as RNAses and inhibit their enzymatic activities.

To be sure that sample quality of tissues shock frozen or conserved in RNAlater for various lengths of time is equally high, we compared the quality of total RNA as well as the expression of 5 different genes including one housekeeping gene (TBP), 3 genes relevant in NSCLC (ERCC1, EGFR, RRM1)(14-18) and one gene (HIF1) regulated by ischemia at least partially on the mRNA level (19-21) in matched lung tumor samples processed by the various procedures. As expected, and has been shown for tissue of other origin (6, 11), we found that the dominant source of variation is interpatient variability. Neither snap freezing nor RNAlater processing for 24 hours or 7 days introduced relevant differences, neither with regard to overall RNA quality nor to the expression of the 5 genes tested.

RNA degradation after withdrawal of tissue from the body may be due to two major reasons. First, the altered environment, especially the lack of oxygen supply, may lead to altered gene expression in the living cells. Second, RNAses may degrade RNA, which, while the cells in the tissue start to die, will not be resynthesized. In both cases, after prolonged storage at room temperature outside of the body, chances in gene expression would be expected. We compared samples that were processed immediately after withdrawal (5 min), or after 20, 60, and 180 min. There was no effect on the measured expression for any of the genes during this time frame. It can be assumed that lack of oxygen does not play a major role for the expression of these genes, and that the cellular structures remain intact for at least up to 3 hours, keeping RNAses compartmented in lysosomes and thus unable to attack the RNA.

Care must be taken not to thaw the samples during shipping or further handling such as preparation of histological slides or isolation of nucleic acids. At this point, another great advantage of RNAlater preservation comes into the play. In a snap frozen tissue sample that thaws, RNA will be degraded immediately after thawing, since freezing/thawing destroys any intracellular compartmentalization and makes RNA accessible to RNAses. In a RNAlater preserved sample protection is maintained after freezing/thawing, and intact RNA with unaltered expression levels can be isolated(7, 9) (and own preliminary data, not shown). Though this is not a licence to interrupt the cold chain, it eases sample handling during analytic procedures, and minimizes temperature effects e.g. during cutting in a microtome cooled to -30°C.

In conclusion, RNAlater is a protective reagent with many in the context of clinical biobanking. As has been shown for tissues of other origin like liver, it is well suitable for conservation of lung tissue prior to long term storage at -80 °C. In comparison to snap freezing in liquid nitrogen, samples show the same quality with regard to overall RNA quality and mRNA expression. Sample handling at different sites of the clinic is easier and safer and enables samples collection even in parts of the clinic not directly connected to a laboratory environment.

